# Crosstalk between Ovate Family Proteins, plant hormones, and microtubule dynamics regulating fruit shape

**DOI:** 10.64898/2026.02.17.706389

**Authors:** Veredas Coleto-Alcudia, Beatriz E. García-Gómez, Christian Dujak, Arnau Fiol, Maria José Aranzana

## Abstract

Fruit shape is a key horticultural trait shaped by conserved genetic pathways and hormonal interactions, yet the mechanisms underlying shape diversity in fleshy fruits remain incompletely understood. Studies in model species such as *Arabidopsis thaliana*, tomato, and rice have established Ovate Family Proteins (OFPs) as central regulators of organ morphology through their interactions with brassinosteroid (BR) and gibberellin (GA) pathways and cytoskeleton dynamics. Here, we combine phylogenetic, transcriptomic, and co-expression network analysis to investigate fruit shape regulation in peach and apple, two major Rosaceae crops. We show that flat and oblong phenotypes are associated with distinct OFP expression patterns and with coordinated changes in hormone-related modules, revealing conserved OFP-hormone-cytoskeleton regulatory circuits. Flat shapes were linked to the activation of flat-associated OFPs in the absence of brassinosteroid signalling, whereas oblong shapes were associated with the activation of elongation-related OFPs under brassinosteroid-responsive conditions. Our findings extend current models of fruit morphology by providing species-specific mechanistic insight into OFP-mediated regulation in Rosaceae, offering a refined framework for breeding fruit shape.

## 1. Introduction

The diversity of fruit shapes observed in nature reflects a complex interplay of genetics, ecological, and evolutionary pressures associated to plant reproduction and seed dispersal mechanisms. In cultivated species, fruit shape variability has arisen through the selection and combination of alleles during domestication and post-domestication events (Liu et al. 2023; Rodríguez et al. 2011; Wu et al. 2018). Today, fruit shape and symmetry are important objectives in fruit breeding due to their strong impact on appearance and quality, which are essential for consumer acceptance and fruit processing. This breeding interest extends beyond fleshy fruits to grains, encompassing fruit trees, vegetables, and arable crops. Recent reviews have emphasized the central role of genetic regulation in shaping fruit morphology across horticultural species (Liu et al. 2024), underscoring the relevance of this trait for crop improvement.

In the Rosaceae family, fruits are highly-valued for their nutritional properties, with species from the *Malus*, *Pyrus*, and *Prunus* genera being among the most widely consumed worldwide. While fruits from *Malus* and *Pyrus* species (apples and pears) are pomes, developed from the enlargement of the receptacle and sepals, those from *Prunus* species (e.g., peach, plum, apricot, cherry) are drupes, formed from the enlargement of the ovary. Despite their anatomical differences, a range of fruit shapes, from elongated to flat, is observed in both pomological types. This diversity is consistent with broader angiosperms patterns showing that fruit morphology is strongly influenced by ovary structure (Xiang et al. 2024). Fruit shape is determined early during floral bud differentiation, followed by a series of molecular and cellular processes throughout development (Mauxion et al. 2021). Gene expression patterns modulate fruit shape through synergistic interactions among developmental regulators and hormonal pathways, influencing traits such as fruit elongation via gynoecium development or post-fertilization growth dynamics (Li et al. 2023).

The diversity of fruit shapes in model and cultivated species has facilitated the study of the genetic mechanisms regulating this trait. However, the economic importance and annual growth habit of crops like tomato and rice have likely driven more intensive research efforts on these species compared to fruit trees. The first gene found to be associated with fruit shape, *OVATE*, was identified in tomato (Ku et al. 1999; Liu et al. 2002). *OVATE* is a member of the Ovate Family Protein (OFP), a protein family unique to land plants, characterized by a conserved C-terminal domain of approximately 70 amino acids (Liu et al. 2014; 2002). Since this discovery, extensive research has focused on understanding the functional roles of this gene family and the regulatory mechanisms that govern organ shape in the model species *Arabidopsis thaliana*, rice, and fleshy fruits (Li et al. 2023; Snouffer et al. 2020).

In *Arabidopsis*, the entire Ovate Family Protein has 20 putative genes (Hackbusch et al. 2005; Liu et al. 2014), which have been described as transcriptional repressors (Wang et al. 2007; 2011) and classified in three classes based on their mutant redundant phenotypes (Wang et al. 2011). In crops such as tomato, rice, and peach, this family encompasses 30, 33, and 16 putative genes, respectively (Huang et al. 2013; Liu et al. 2014; Yu et al. 2015). In general, the overexpression of OFPs results in the development of shorter but wider organs, as *SlOFP20* in tomato (Zhou et al. 2019), *AtOFP1-4* in *Arabidopsis* (Li et al. 2011; Wang et al. 2011), *OsOFP13* (annotated as *OsOFP19* in (Yu et al. 2015)) in rice (Yang et al. 2018), and *PpOFP1* in peach (Zhou et al. 2021). In apple, a GWAS analysis highlighted *MdOFP4*, in Chromosome 11, as candidate for the ration between fruit elongation and width (also known as Fruit Shape Index, FSI) (Dujak et al. 2024). However, other OFP genes, such as the *OsOFP1/10/30* genes in rice (annotated as *OsOFP1/14/8* in (Yu et al. 2015)) produce elongated grains (Zhao et al. 2018), suggesting different pathways of the OFPs in organ shape regulation.

Plant hormones regulate cell proliferation and expansion during fruit development and growth. In tomatoes, strong connections have been established between fruit morphology and hormones such as auxins, gibberellins (GAs), brassinosteroids (BRs), and ethylene. The application of exogenous auxins in tomato results in ovary elongation and modifies fruit cell size and number (Wang et al. 2019). In peach, levels of indole-3-acetic acid (IAA), the main endogenous auxin, were significantly higher in round than in flat fruits, and four genes in the auxin signaling pathway were proposed to be involved in flat fruit shape determination, pointing to a potential interaction between *PpOFP1* and auxin signaling (Guo et al. 2018). However, studies in tomato indicate that auxin is not directly involved in the pathway mediated by the *OVATE* gene (Wang et al. 2019).

Brassinosteroids are plant steroid hormones that bind to the BRASSINOSTEROID INSENSITIVE 1 (*BRI1*) receptor in the cell membrane and activate a phosphorylation cascade. This cascade culminates in the dephosphorylation and subsequent activation of the transcription factors BRASSINAZOLE-RESISTANT 1 (*BZR1*) and BRI1-EMS-supressor 1 (*BES1*) by PROTEIN PHOSPHATASE 2A (*PP2A*) (Wang et al. 2002; Yin et al. 2002). When these factors are active, they enter in the nuclei and promote the transcription of BR-related genes (Vert and Chory 2006; Zhao et al. 2002). In addition, *PP2A* facilitates *BRI1* turnover when is methylated and activated by SUPPRESSOR OF BRI1 (*SBI1*) (Wu et al. 2011). In the absence of BRs, the BRASSINOSTEROIDS INSENSITIVE 2 (*BIN2*) kinase phosphorylates *BZR1*/*BES1*, inactivating the BRs cascade (He et al. 2002; Sun et al. 2010; Yin et al. 2002).

Several studies have established a role for OFPs in the BR response in rice, while their function in tomato fruit shape is largely unknown (Li et al. 2023). *OsOFP1*, *OsOFP2*, and *OsOFP30* (*OsOFP1*, *OsOFP3* and *OsOFP8* in (Yu et al. 2015) interact with the homolog of *BIN2* (Xiao et al. 2017; 2020; Yang et al. 2016), while *OsOFP13* (*OsOFP19* in (Yu et al. 2015)) activates BR catabolic genes, resulting in both cases in the BR signaling cascade inactivation. Besides, plants overexpressing *OsOFP30* (*OsOFP8* in (Yu et al. 2015)) show reduced expression levels of BR synthesis genes (Yang et al. 2016).

A link between BRs and the cytoskeleton has been described in tomato, where the overexpression of a *BZR1* homolog, *SlBZR1.7*, promotes fruit elongation by the upregulation of *SUN* gene (Yu et al. 2022), a member of the IQ67 domain calmodulin-binding gene family associated with microtubules and involved in cytoskeleton organization (Bürstenbinder et al. 2017). In addition, the PP2A protein, involved in the BRs cascade, controls microtubule dynamics through its subunit *TON2*/*FASS* (*TONNEAU2*), as well as cell division forming a TTP complex with *TON1* (*TONNEAU1*) and *TRM* (TONNEAU1 Recruiting Motif) (Spinner et al. 2013; Yoon et al. 2018). In tomato, it has been described the interaction of *OVATE* and *SlOFP20* with TRM proteins, regulating their subcellular localization to control cell division (Wu et al. 2018) and organ shape. Recently, Li et al. (2023) reviewed the studies in fruit shape regulation, focusing on the Ovate Family Protein-*TONNEAU1* Recruiting Motif (OFP-TRM) and IQ67 domain (*IQD*) regulatory pathway. They concluded that, while significant progress has been made in understanding fruit shape regulation in Arabidopsis and rice, there has been limited exploration of these mechanisms in fleshy fruits.

The OFPs are also involved in the regulation of other hormones. For example, the *AtOFP1* protein represses *AtGA20ox1* (Ding et al. 2020; Wang et al. 2007), a gibberellin synthesis gene producing shorter organ cells, suggesting a connection between OFPs and GAs in the control of organ shape. In addition, the overexpression of *SlOFP20* revealed a possible crosstalk between BR and GA in tomato (Zhou et al. 2019), which is synergistic to *OVATE* in shape regulation by primarily modulating cell division patterns (Wu et al. 2018).

While these and other works suggest a crosstalk between OFPs, BRs as well as other hormones, and the cytoskeleton, determining organ shape, a holistic view of the mechanism in fleshy fruits is still missing. Here we used transcriptomics and phylogenetics and constructed a gene co-expression network to understand the regulation of shape in fruits of the Rosaceae family (apple and peach), with an expected impact in enhancing fruit breeding.

## 2. Results

### 2.1. Phylogenetic analysis of OFPs

The alignment of the entire amino acid sequences of 127 OFPs from Arabidopsis (20), tomato (30), rice (33), apple (28), and peach (16) revealed that the OVATE domain was highly conserved in all but 14 of them, which lacked it (Figure S1). This high conservation contrasted with the lower conservation level on the rest of the amino acid sequence. A phylogenetic analysis produced a tree with nine clades (Figure 1). The OFPs with a described role in the flat shape, such as PpOFP1, AtOFP1/2/4, SlOFP20, and OsOFP4/13, clustered in clade 6 and 7 (blues), while the OFPs known to be associated with elongated rice grains, such as OsOFP10, OsOFP1 and OsOFP30, were in clades 1 (purple) and 9 (orange). The proteins without or with a short OVATE domain clustered in clades 3 and 4 (green).

**Figure 1.**
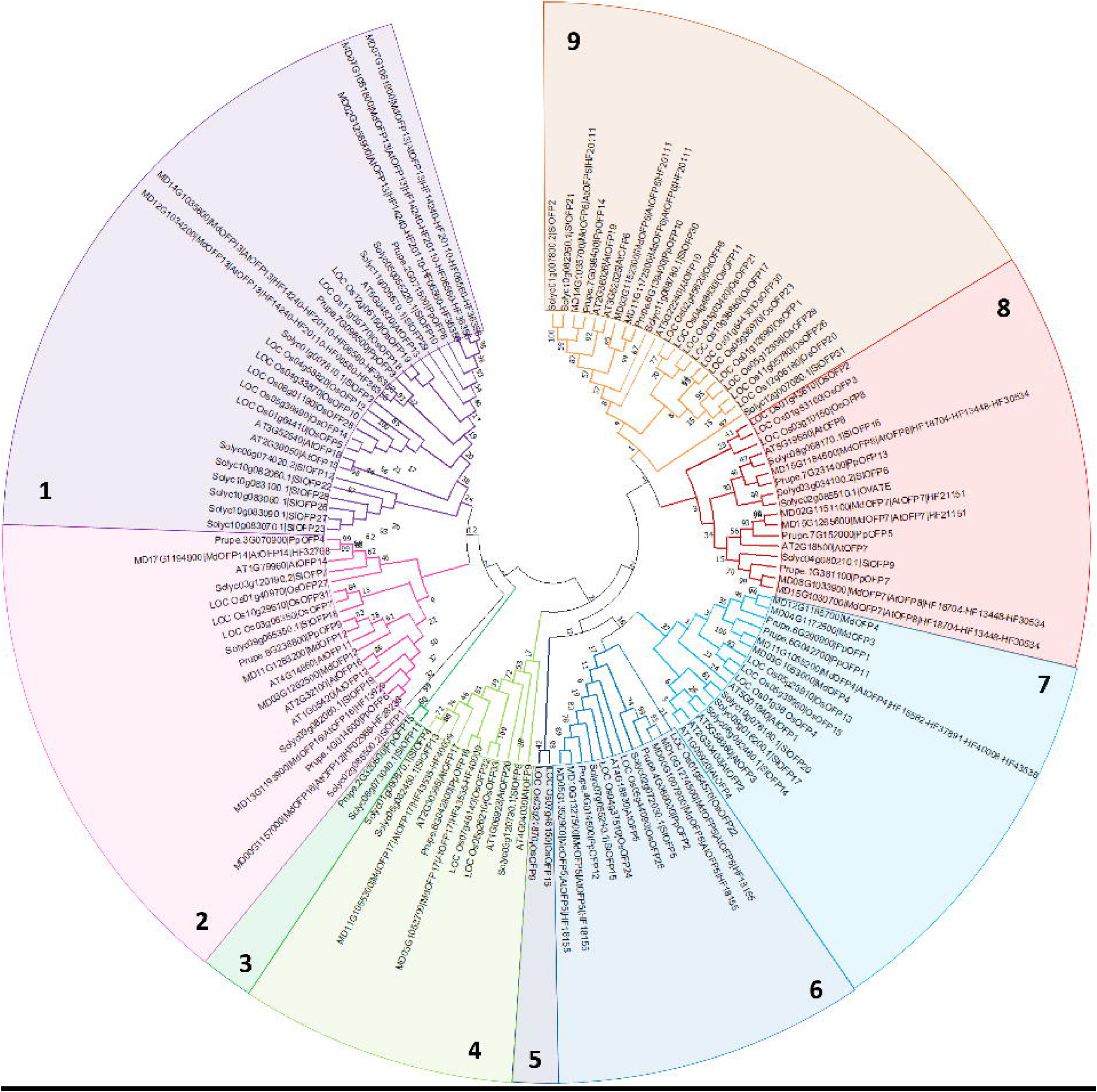
Phylogenetic analysis of Ovate Family Proteins (OFPs) across five plant species. Maximum Likelihood phylogenetic tree of 127 OFP proteins from *Arabidopsis thaliana*, *Solanum lycopersicum*, *Oryza sativa*, *Prunus persica*, and *Malus domestica*. Distinct clades are indicated by numerical labels and color coding.

Within the amino acid sequences, we identified five conserved motives. Motives 1 and 2 (Figure S2) were present in most of the OFPs and corresponded to the C-terminal domain characteristic to the Ovate Family Proteins (Interpro ID IPR006458, Pfam ID PF04844). Motif 3, annotated as a DNA binding domain often found in OFPs (Interpro ID IPR025830, Pfam ID PF13724), was only observed in those associated with flat shape (clades 6 and 7 of the phylogenetic tree). Motif 4 was found in eight out of the 26 OFPs of clade 1, with OFPs associated with elongated grains. Motif 5 was largely found within the sequences of the clade 2. No Interpro/Pfam domain annotations were found for motives 4 and 5.

### 2.2. Differential gene expression analysis between fruit shapes

To evaluate the functional conservation of the fruit shape related OFP genes between Rosaceae species, we conducted a whole genome gene expression analysis (RNAseq) in flower buds and in early developed fruits of a flat peach variety (UFO) and of its round somatic mutant (MUT) (Figure 2A), and in apple fruits of three varieties at three stages of development (13, 61 and 98 days after anthesis (DAA)) (Figure 3A). These varieties showed contrasting fruit shape: Grand’mere’ (Gra) with flat shape, ‘Kansas Queen’ (Kan) round, and ‘Skovfoged’ (Sko) oblong.

**Figure 2.**
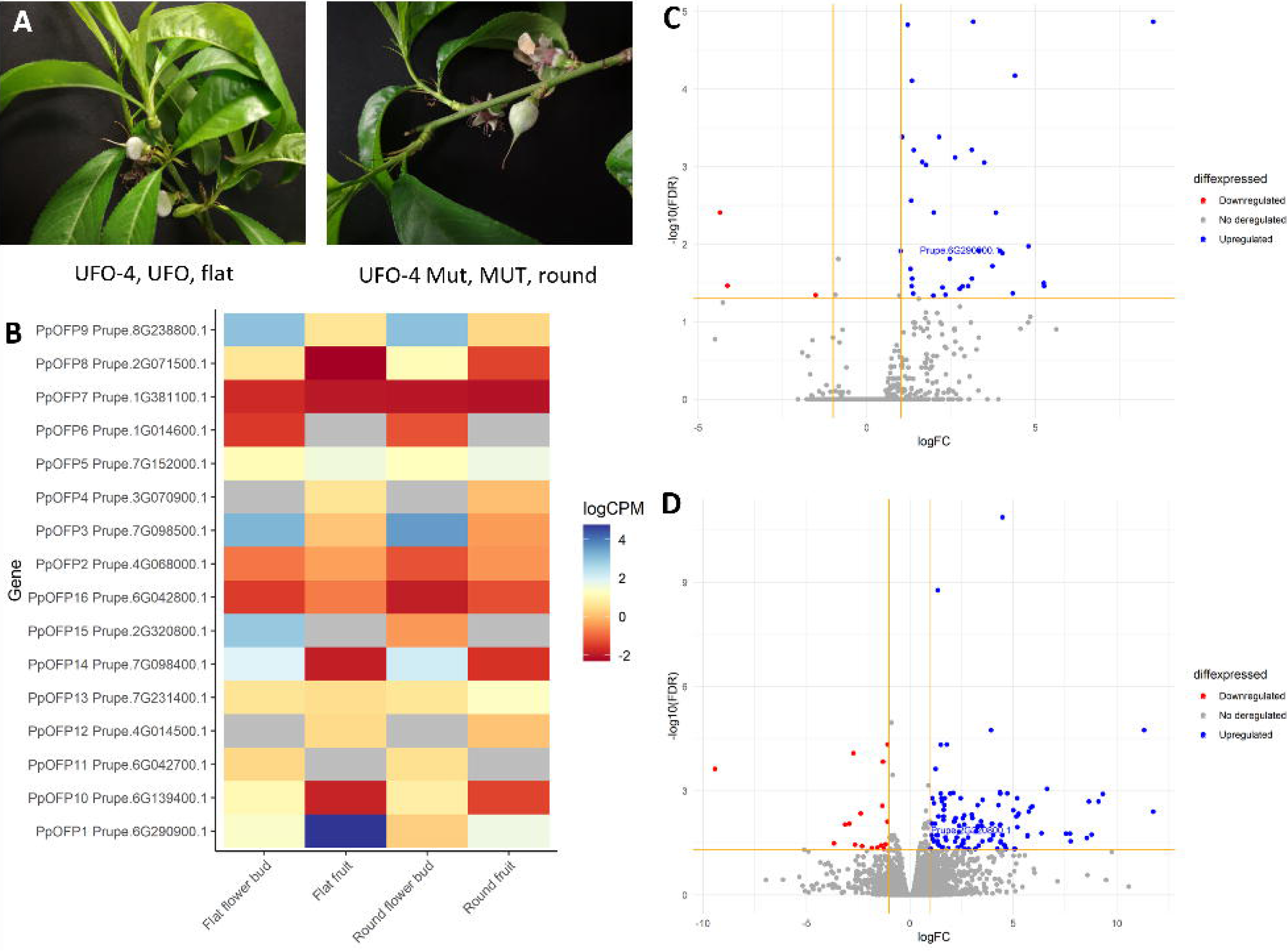
Transcriptomic analysis of fruit shape and developmental stages in *Prunus persica*. (A) Representative images of UFO and MUT peach fruits used for transcriptome profiling. (B) Expression patterns of PpOFPs genes across fruit shape and developmental stages. Colors indicate logCPM values; grey denotes undetected expression. (C-D) Vulcano plots of DEGs (FDR<0.05, |logFC|>1) in UFO *vs* MUT pairwise comparisons for (C) fruit and (D) flower buds. Red= downregulated genes, blue=upregulated genes. Deregulated PpOFPs are labelled.

**Figure 3.**
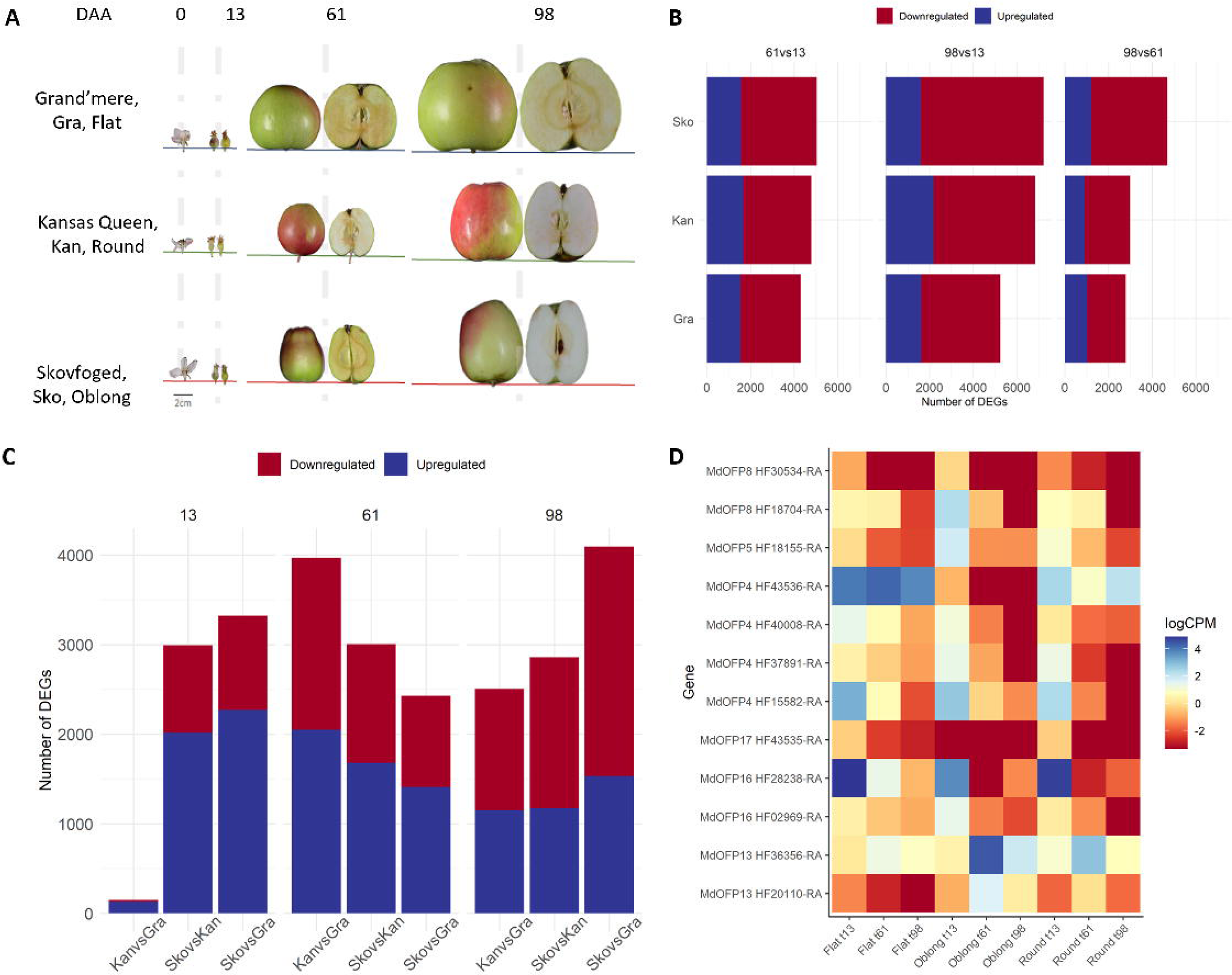
Transcriptomic analysis of fruit shape and developmental stages of *Malus domestica*. (A) Representative images of apple fruits used for transcriptome profiling. (B) Number of DEGs identified across developmental stages within each variety. (C) Number of DEGs identified across varieties at each developmental stage. Red = downregulated, blue = upregulated. (D) Expression pattern pf MdOFPs across fruit shapes and developmental stages. Colors indicate logCPM values; grey denotes undetected expression.

Principal Component Analysis (PCA) of peach transcriptomes revealed that gene expression patterns clustered strongly by developmental stages (flower buds vs. fruit) while no clear separation was observed between UFO and MUT (Figure S3A). Consequently, pairwise comparisons between the UFO and MUT samples were performed independently within each stage. A higher number of differentially expressed genes (DEGs) were found in flower buds (185) compared to fruits (45) (Figure S3B, Table S3) with only two genes’ transcripts (Prupe.4G171800.1, a NAC domain containing protein; and Prupe.5G034800.1, a Glycine-rich protein family) shared between the two stages. None of these genes have been previously associated with fruit shape development. Only two OFP genes were found to be differentially expressed in either flower buds or fruits (Figure 2B-D). The gene PpOFP15 (Prupe.2G320800), which clustered within the genes without the OVATE domain in the phylogenetic tree, was upregulated in the flower buds of the flat UFO genotype, while PpOFP1 (Prupe.6G290900) was upregulated in the flat fruits. As can be seen in the vulcano plots (Figure 2C-D), the majority of the DEGs were predominantly upregulated in the UFO (flat) buds and fruits compared to the round mutant.

In apple, the PCA revealed distinct transcriptomic profiles primarily driven by developmental stage, with additional variation associated with fruit morphology. The first two principal components clearly separated samples according to developmental timing (Figure S4A), while differences related to fruit shape became apparent after plotting components two and three (Figure S4B). Thus, pairwise comparisons of RNA-seq data were performed across developmental stages (98 *vs* 13, 61 *vs* 13 and 98 *vs* 61) and between varieties at each stage (Sko *vs* Gra, Sko *vs* Kan and Kan *vs* Gra) (Figure 3B-C, Figure S4C-E, Table S4). The largest number of DEGs was observed between the earliest and the latest developmental stages (13 *vs* 98 DAA), totaling 3,136 DEGs (Figure S4C), reflecting major reprogramming during fruit growth and maturation. The comparison between 61 and 13 DAA revealed 1,760 DEGs, consistent with active cell division and peak expansion. In contrast, the fewest DEGs (702) were found between 61 and 98 DAA, corresponding to the transition toward ripening, when transcriptional activity stabilizes, and cell expansion slows.

The number of DEGs identified in pairwise comparisons between apple varieties at each developmental stage varied substantially. The contrast between varieties with extreme fruit shapes (Skov *vs* Gra) yielded the highest number of shared DEGs across stages (957), whereas the comparison between flat and round fruits (Kan *vs* Gra) produced the lowest number (108) (Figure S4D). Across all variety pairs, the majority of shared DEGs were detected at 61 DAA (328), whereas only nine genes were consistently differentially expressed in all varieties at 13 DAA (Figure S4E).

The GO and KEGG enrichment analysis of the DEGs between different scenarios identified a large fraction of genes involved in cell division (e.g., DNA packaging complex, DNA binding, nucleosome) and microtubule movement (e.g., microtubule binding, microtubule-based movement, microtubule motor activity) at the early stages of development (Figure S5 A-B, D-E, G-H), while at later stages the enriched genes were related to cell growth, expansion and elongation (e.g., growth factor activity, nucleosome organization) (Figure S5 C, F, I). Also, at the later stages, auxins, cytokinins, brassinosteroids, gibberellins, jasmonic acid and salicylic acid genes were downregulated, and abscisic acid and ethylene genes were upregulated (Figure S6). In the pairwise comparisons of varieties at different developmental stages (Figure S7), the gene ontology enrichment analysis (GOEA) identified transcription-related and kinase activity terms (e.g., DNA-binding transcription factor activity, sequence specific DNA-binding, protein kinase activity, MAPK signaling pathway) at 13 and 61 DAA (Figure S7A-B, D-E, G-H), as well as terms related to size and cytoskeleton organization through actin (F-actin capping protein complex) at 98 DAA (Figure S7C, F, I). KEGG enrichment analysis highlighted hormonal changes during fruit development and ripening. The brassinosteroid synthesis pathway showed differential regulation across fruit shapes, with upregulation observed in oblong apples at the early developmental stages (Figure S7G).

Examining the expression patterns of OFPs, we observed a general downregulation during the later developmental stages compared to the earlier stages (Figure 3D). Additionally, regardless of the developmental stage, HF43536 (MdOFP4) was upregulated in flat apples, while HF20110 (MdOFP13) was downregulated (Figure 3D). MdOFP4 clustered in clade 7 of the phylogenetic tree (Figure 1) with other OFPs related to flat shapes as PpOFP1, AtOFP1/2/4 and SlOFP20, while MdOFP13 with OFPs related to cylindrical shapes as OsOFP10 in clade 1.

### 2.3. Identification and characterization of gene modules associated with different fruit shapes

A weighted gene co-expression network analysis (WGCNA) was conducted to identify and elucidate the connections between gene expression patterns. For that purpose, we used the unique genes identified from the RNAseq data in peach fruits (20,507), flower buds (21,856), and in apple (25,565).

In peach fruits, the WGCNA grouped the genes in 105 modules (Figure 4A, Figure S8A-B). Each module was tested for its association with fruit shapes. As shown in Figure 4A, six of them (‘bisque4’, ‘darkseagreen1’, ‘lightcyan1’, ‘antiquewhite4’, ‘mistyrose’ and ‘palevioletred’) were significantly associated with the round or flat shapes (*p*-value <0.05). These modules contained 29 out of the 45 DEGs mentioned earlier (Table S5), being PpOFP1 (upregulated in flat peaches) in the ‘palevioletred’ module, which also contained four additional DEGs. Hierarchical clustering of the 193 genes within this co-expression module (Figure S8C-D) revealed nine distinct clusters. PpOFP1 was found in cluster 4. The other four DEGs (a plant invertase/pectin methylesterase inhibitor in chromosome 1, two Frigida-like proteins in chromosome 6, and a glyoxal oxidase-related protein) were found in cluster 3 together with other downregulated genes in flat fruits. Cluster 8 contained genes that were upregulated in flat fruits.

**Figure 4.**
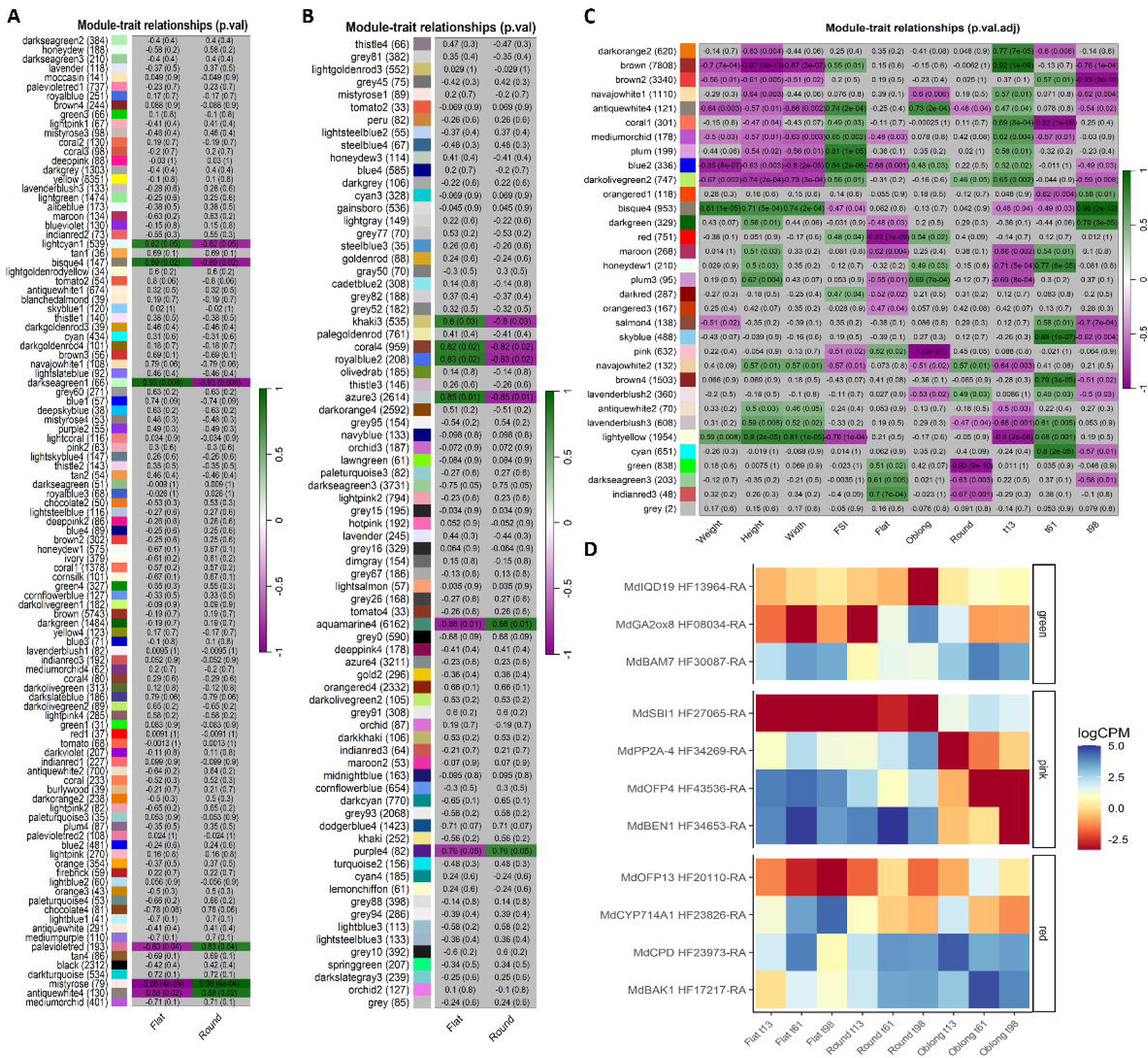
Weighted Gene Co-expression Network Analysis (WGCNA) of transcriptomic data. (A-C) Correlation matrices between co-expressed modules (y-axis) and fruit shapes (x-axis) in peach fruits (A), peach flower buds (B), and apple fruits (C). Cell color indicates correlation coefficient. (D) Expression patterns of DEGs associated with organ shape in apple “pink”, “red” and “green” modules. Colors indicate logCPM values.

Following the same approach in the flower bud RNA-seq dataset, six modules (‘aquamarine4’, ‘azure3’, ‘coral4’, ‘khaki3’, ‘purple4’ and ‘royalblue2’) out of the 77 identified by the WGCNA correlated with fruit shape (Figure 4B). These modules comprised 95 out of the 185 DEGs identified (Table S6). The gene PpOFP15 (Prupe.2G320800), which was upregulated in the flower buds of the UFO genotype, grouped in cluster 9 of the ‘aquamarine4’ module. In this module, clusters 7, 8 and 9 contained genes upregulated in flower buds of the UFO, while cluster 6 contained those that were downregulated (Figure S9C-D).

In the apple dataset, correlation analysis between gene modules and traits showed that the ‘pink’, ‘red’, ‘green’, ‘darkred’, ‘orangered3’ and ‘indianred3’ modules correlated with fruit shape, defined by shape classification (flat, round, and oblong) or by the shape ratio FSI, rather than size-related parameters or developmental stages (Figure 4C). These modules collectively contained 1,829 DEGs. Hierarchical clustering of genes within the modules with the strongest correlation (‘pink’, ‘red’ and ‘green’) (Figure 4C), identified clusters with different regulation patterns (Figure S10, Table S7). Genes in the ‘pink’ module were differentially regulated in oblong fruits (Figure S9C), in the ‘red’ module in flat fruits (Figure S9D), and in the ‘green’ module in round fruits (Figure S10E). Brassinosteroid and OFP genes were identified in the ‘pink’ (e.g. AtOFP4/MdOFP4, MdBEN1 in cluster 6, MdPP2A in cluster 1, and MdSBI1 in cluster 2) and ‘red’ (e.g. AtOFP13/MdOFP13 in cluster 2; MdCPD and MdBAK1 in cluster 5) module, while the ‘green’ module contained genes related to microtubule dynamics (e.g. MdIQD19 in cluster 4) and gibberellins (e.g. MdGA2ox8 in cluster 5) (Figure 4D).

## 3. Discussion

Members of the Ovate Family Protein have been consistently described as regulators of seed and fruit shape, either flat or elongated, suggesting that the mechanism of shape variation may be highly conserved between species and that the OFP genes are key players. Here we confirmed that the OVATE domain was highly conserved among OFPs of different species, while considerable differences were observed between the non-OVATE domains. It is particularly interesting the conservation of a DNA binding domain in the OFP protein sequences of the genes with a described role in flat shape determination, while this domain was not present in the OFPs reported to be related to cylindrical shape. This may suggest that the OFPs inducing flat shape could execute their function by directly binding to the DNA, while the cylindrical shape-related OFPs could interfere with the transcriptional machinery by binding to other partners.

To investigate a possible crosstalk between OFPs, genes regulating the cytoskeleton dynamics, BRs and GAs hormones in Rosaceae fruits, we performed a transcriptomic analysis in peach with flat and round shape and in three apple varieties described as flat, round and oblong.

The PpOFP1 was found upregulated in flat peach fruits, which was in agreement with (Zhou et al. 2021), who found PpOFP1 overexpressed in flat peaches. PpOFP1 interacts and sequesters PpTRM17, a TONNEAU1 Recruiting Motif (TRM) protein involved in fruit elongation in tomato (Zhang et al. 2023), through its interaction with OFPs; probably in conjunction with IQ67 Domain (IQP) proteins (Snouffer et al. 2020). The gene co-expression network analysis revealed that the PpOFP1 gene occurred in the same module as PpIQD26 gene, a member of the IQ67 calmodulin-binding family protein, which is consistent with the possible link between OFPs and microtubule genes in shape regulation suggested by Snouffer et al. (2020).

In addition to Prupe.6G290900 (PpOFP1), other genes on chromosome 6 of fruits were differentially expressed between the UFO and its sport mutant (MUT). This mutant was generated after a spontaneous mutation in one meristematic bud, which gave place to a branch producing round fruits (López-Girona et al. 2017). Studies on these materials revealed a loss of heterozygosity (LOH) at the distal end of chromosome 6 extending a 6.5 Mbp, resembling a non-homologous recombination after a chromosome break which ended in the replacement of the region containing the flat-associated allele with the homologous chromosome, which contained the variant producing round fruits. Thus, while the original tree was heterozygous for the flat and round-associated alleles, the mutant was homozygous for the round one. Due to the size of the chromosome region replaced, other genes in phase with the flat-associated allele were also substituted and, therefore, their transcripts should appear differentially regulated between the UFO and the MUT fruits. This may apply to five genes located downstream PpOFP1. Also, we identified four genes at the proximal end of chromosome 6 that are overexpressed in flat fruits. Given that flat and round fruits share an identical DNA sequence in this region, it is plausible that their differential expression is regulated by genes located at the distal end of chromosome 6. One of this genes at the proximal end of chromosome 6 is the Prupe.6G016700, a receptor like protein kinase associated with fruit polar diameter by (Elsadr 2016), and co-localizes with a QTL for fruit shape and size identified in a collection of non-flat peaches by (Cirilli et al. 2021).

Interestingly, the expression level of the PpOFP1 gene was only marginally higher in the buds of the flat genotype, without reaching statistically significant differences. Instead, in flower buds, the PpOFP15 was found significantly upregulated. However, the lack of OVATE domain in the protein sequence of PpOFP15 suggests that it may not be involved in shape regulation. Given that fruit shape is already determined at the early stages of flower development, as clearly observed in the ovaries (López-Girona et al. 2017), this data suggest that fruit shape determination may actually occur at stages shortly after the stage C of the Baggiolini scale.

The gene co-expression network grouped the PpOFP3 gene together with PpOFP1 and PpOFP15 in the ‘aquamarine4’ module, although following an opposite expression pattern. This aligns with reports in other species where distinct OFP members regulate the development of different shapes (Wang et al. 2011; Zhao et al. 2018; Zhou et al. 2021). Besides these OFPs in the ‘aquamarine4’ module, several genes linked to fruit shape were identified, including PpIQD23, PpCYP90D1, PpBSK1, PpGA3ox1, and PpGA2ox2. PpIQD23, a member of the IQ67 calmodulin-binding protein family, is known for its association with elongated fruit shape in tomatoes (Yu et al. 2022). This gene family is connected to microtubules through the TRM/OFP/IQD pathway (Bao et al. 2022; Huang et al. 2013; Lazzaro et al. 2018; Snouffer et al. 2020). PpCYP90D1 is associated with BR synthesis, PpBSK1 with BR signaling, PpGA3ox1 with GA synthesis, and PpGA2ox2 with GA inactivation (Wang et al. 2007). The presence of these hormone-related genes co-expressed with PpOFPs in the same module suggests that fruit shape regulation in peach involves both microtubule interactions and hormone signaling (Chen et al. 2021; Li et al. 2023; Snouffer et al. 2020).

The transcriptome of fruits of three apple varieties of different shapes (flat, round, and oblong) at three developmental stages enabled a deeper exploration of fruit shape determination across development. Although the varieties studied differed in shape as well as in many other traits, the larger number of DEGs between extreme shapes suggests that a significant number of these differences could be caused by fruit shape. The earliest stages of fruit development showed higher regulatory activity than the latest, as could be concluded by the larger number of DEGs at the initial stages. GO and KEGG enrichment analysis of differentially expressed genes across developmental time points for each shape showed that early development stages are characterized by cell division, followed by cell expansion and elongation in later stages in consistence with (Janssen et al. 2008). The regulation of fruit development is largely influenced by hormones, as well as by their interactions (McAtee et al. 2013; Simonini and Østergaard 2019; Garrido-Auñón et al. 2024; Liu et al. 2024; Rafiq et al. 2025). Following flower fertilization, auxin is produced by the developing seeds and promotes gibberellin acid (GA) biosynthesis (Devoghalaere et al. 2012; Serrani et al. 2007), so fruit set is mainly mediated by auxins, GAs, cytokinins (CKs) and brassinosteroids (BRs) (Fu et al. 2008; Mariotti et al. 2011). As the cell division phase comes to an end, fruit growth is induced by auxins, GAs and BRs (Davies 2010; Devoghalaere et al. 2012; Pattison and Catalá 2012). During fruit maturation and ripening, the primary hormones responsible are auxin in combination with CK, and ethylene in combination with ABA, respectively (Giovannoni 2007; McAtee et al. 2013). Consistently, our findings revealed that auxins, cytokinins, brassinosteroids, gibberellins, jasmonic acids, and salicylic acid are downregulated at the later stages of fruit development, while abscisic acid and ethylene are upregulated. To overcome the complexity of the apple transcriptomic analysis, a Weighted Gene Correlation Network Analysis (WGCNA) was performed using all genes, and not just DEGs, to decipher the regulatory networks controlling fruit shape. Three modules (pink’, ‘red’ and ‘green’) were highly correlated with the fruit shape index (FSI) and the different shapes, but not with developmental stages, so we retained those modules for a detailed study of fruit shape regulation.

In the ‘pink’ module, MdOFP4, MdBEN1 and MdPP2A-4 were downregulated in oblong apples (Figure S4 C-E), while MdSBI1 was upregulated. According to the phylogenetic analysis, MdOFP4 is the homolog of AtOFP1-4, PpOFP1, SlOFP20, OsOFP6/19/15, all related to flat organ shapes (Chen et al. 2021; Ma et al. 2022; Sun et al. 2020; Wang et al. 2007; Xiao et al. 2017; Yang et al. 2018; Zhou et al. 2021; 2019). Therefore, the downregulation of this gene in oblong apples is consistent with previous literature where a GWAS revealed significant association between MdOPF4 expression and fruit shape (Dujak Riquelme 2023; Dujak et al. 2024), suggesting that this protein is conserved not only in its structure, as demonstrated in the phylogeny analysis, but also in its functional role, possibly being a negative regulator of elongation. Interestingly, the other three genes are related to the brassinosteroid synthesis and signalling. BEN1 is involved in the brassinosteroid catabolism (Yuan et al. 2007), and SBI1 expression is activated upon BR signaling, causing the accumulation of BR-activated BRI1 and the methylation of PP2A (Wang et al. 2012; Wu et al. 2011). PP2A has a dual activity in BR signaling due to its involvement in the BRI1 turnover and the BZR1 dephosphorylation to activate BR response (Wang et al. 2012; Wu et al. 2011). In addition, PP2A-4, a member of subfamily II, is involved in microtubule organization through its interaction with TON2. This interaction influences cell shape and, subsequently, organ shape (Yoon et al. 2018; Zhang et al. 2020). The co-expression of these genes found in oblong apples suggests that there is a tight regulation of fruit shape through BRs and OFPs: in oblong apples, BR synthesis is active by the downregulation of BR catabolism through BEN1, and the BR signalling is active by the upregulation of SBI1. The downregulation of PP2A in oblong apples provides a fine-tuned homeostasis of BR signalling while ensuring the good organization of microtubules because TON2 is not able to interact with PP2A. Elongation, mediated by microtubules organization, is also affected by the downregulation of MdOFP4. Reduced expression of OFPs leads to the mis-localization of TRMs and results in elongated organs. Additionally, MdOFP4 is homologous to AtOFP2, whose expression is reduced by BR treatment (Zhang et al. 2020). It is also related to SlOFP20 and OsOFP13, both of which are negative regulators of BR response. Overexpression of these genes leads to increased levels of BR catabolism genes (Yang et al. 2018; Zhou et al. 2019). To sum up, ‘pink’ module analysis suggests that in oblong apples, BR signalling produce elongated shape through the inactivation of MdOFP4 and the consequent organization of microtubules through TRMs.

In the ‘red’ module, the genes MdOFP13, MdBAK1, MdBSK3, MdBSK7, MdCPD and MdCLE1 were downregulated in flat apples, while MdBSK1 and MdCYP714A1 were upregulated. MdOFP13 is the homologous of OsOFP10 (Figure 1), related to elongated rice grains by inhibiting GS9 (Zhao et al. 2018). Therefore, its downregulation in flat apples is in concordance with the literature. Also, the downregulation of the BR signaling genes BAK1, BSK3 and BSK7 in flat apples suggests that the BR response is inactivated in flat apples, what matches with the additional downregulation of CPD, a BR biosynthesis gene. However, BSK1 is found to be upregulated in flat apples, probably as a compensatory effect of BR signaling or related to immunity because both BSK7 and FLS2, immunity-related pathway, were also downregulated in flat apples. Interestingly, CYP714A1, a gibberellin (GA) inactivation enzyme that is associated with dwarf shorter plants (Zhang et al. 2011), was upregulated in flat apples. This ‘red’ module analysis suggests that the inactivation of MdOFP13, and BR and GA signaling avoid the organization of microtubules, resulting in the flat shape of apples.

In the ‘green’ module, the genes MdIQD19 and MdBAM7 were downregulated in round apples, while MdGA2ox8 was upregulated. Interestingly, IQD19 is a member of the IQ67 calmodulin-binding family protein, as well as the PpIQD26 gene found in the co-expression network analysis in peach commented above, and was associated with elongated shape in tomato (Yu et al. 2022) through TRM/OFP/IQD-related microtubule dynamics (Bao et al. 2022; Huang et al. 2013; Lazzaro et al. 2018; Snouffer et al. 2020). BAM7 together with BAM8 are the two members of the beta-amylase family proteins that contains BZR1-type DNA binding domains (Monroe and Storm 2018; Reinhold et al. 2011; Soyk et al. 2014). The downregulation of both IQD19 and BAM7 in round apples suggest a possible role of BZR1-BAM genes in modulating the transcription of IQ67 proteins, such as IQD19, similarly to the regulation found in tomato through the SUN gene (Yu et al. 2022). GA2ox8 is a GA inactivation enzyme, with the same role as CYP714A1 found to be upregulated in flat apples, although with the opposite role than GA20ox and GA3ox (Hedden and Phillips 2000). Interestingly, it is known that AtOFP1 represses AtGA20ox1, inhibiting GA synthesis (Wang et al. 2007). Although no OFP has been found in this ‘green’ module, the fact that GAs are inactive supports the hypothesis that fruit shape is regulated by OFPs through an interaction between BRs and GAs.

In conclusion, this research on the regulation of fruit shape in the Rosaceae family, with a focus on peach and apple, provides valuable insights into the molecular mechanisms governing fruit morphology, unveiling key regulatory elements. The phylogenetic, transcriptomic and regulatory network analyses carried out in this study revealed that flat shapes are regulated by the activation of flat-related OFPs, such as PpOFP1 and MdOFP4, in absence of brassinosteroids, while oblong shapes are regulated by the activation of oblong-related OFPs, as MdOFP13, that drives to an organization of microtubules in presence of brassinosteroids (Figure 5). These results align with findings in model species such as Arabidopsis, rice, and tomato, indicating potential conservation in fruit shape regulation across diverse plant species. Phylogenetic analysis confirmed the sequence conservation of OFP protein sequences in apple and peach, highlighting the evolutionary relationships within the Rosaceae family. Overall, our findings suggest an intricate regulatory network involving OFPs, cytoskeleton dynamics, brassinosteroids, gibberellins, and other hormonal pathways in shaping fleshy fruits. Understanding these regulatory mechanisms not only enhances our knowledge of fruit development but also provides valuable insights for crop breeding efforts. Further research into the specific interactions and signaling pathways will contribute to unraveling the complex web of factors influencing fruit morphology in the Rosaceae family.

**Figure 5.**
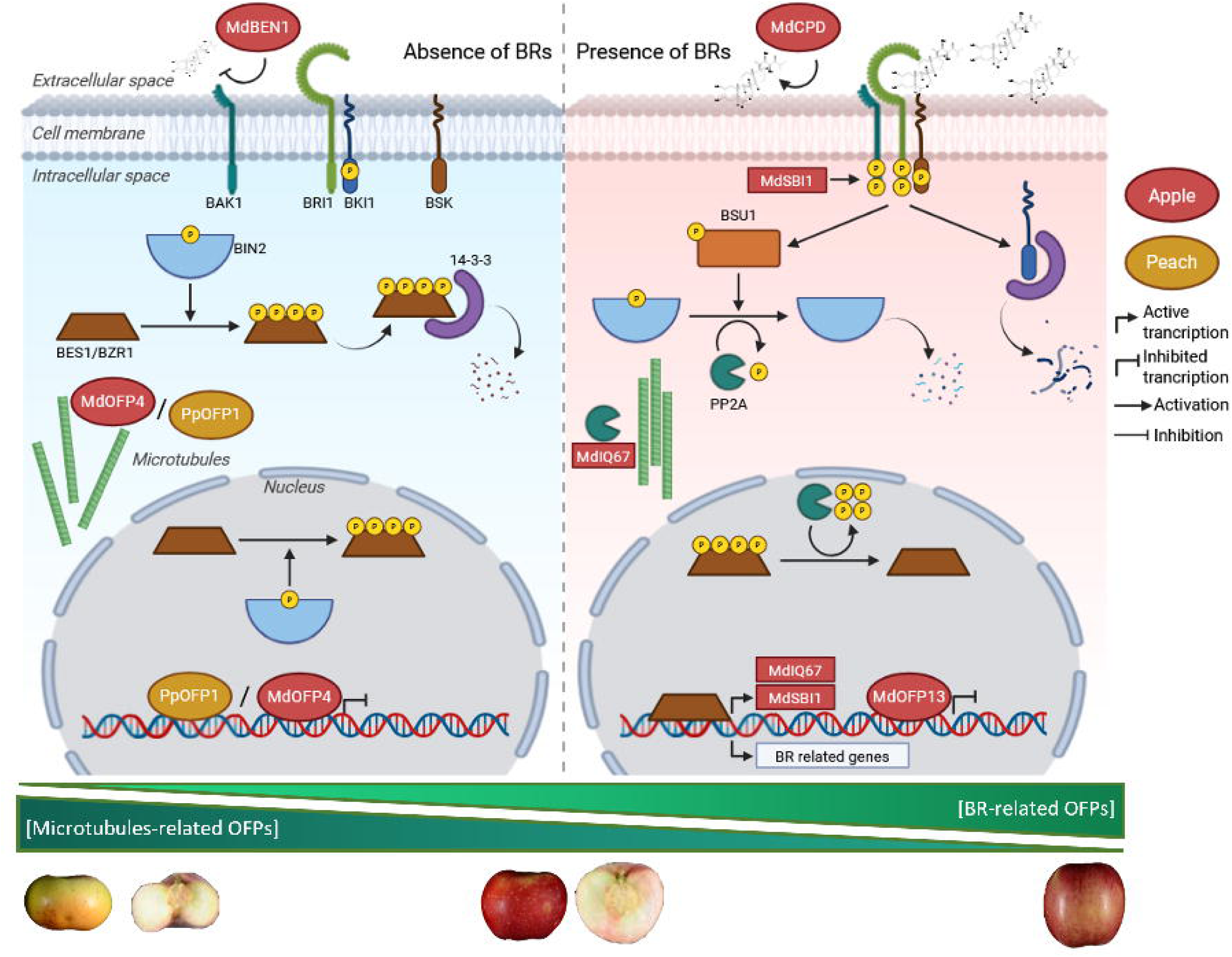
Regulatory model of fruit shape determination in Rosaceae. In the absence of brassinosteroids (BRs), OFPs linked to microtubule dynamics are active, regulating flat fruit shapes. In the presence of BRs, OFPs associated with oblong shapes are active and IQ67 proteins organize microtubules. Figure created with Biorender.

## 4. Materials and methods

### 4.1. Phylogenetic and motif analysis of OFP sequences

One hundred twenty-seven OFP amino acid sequences retrieved from the literature were used in the analysis: 20 from *Arabidopsis thaliana* (Liu et al. 2014), 30 from *Solanum lycopersicum* (Huang et al. 2013; Liu et al. 2014), 33 from *Oryza sativa* (Liu et al. 2014), 28 from *Malus domestica* (Li, Dong, Zhao, et al. 2019), and 16 from *Prunus persica* (Li, Dong, Zhu, et al. 2019; Liu et al. 2014) (Table S1). We used the OFP nomenclature of (Liu et al. 2014) for *Arabidopsis thaliana*, *Prunus persica* and *Oryza sativa*; the nomenclature of (Huang et al. 2013) for *Solanum lycopersicum*; and the nomenclature of (Li, Dong, Zhao, et al. 2019) for *Malus domestica*.

MEGA11 (Tamura et al. 2021) was used for the alignment of the 127 OFP sequences with Muscle algorithm, 50 max iterations and default parameters. Phylogeny tree was constructed with the Maximum Likelihood method, bootstrap test with 1000 replicates and the Jones-Taylor-Thornton (JTT) model of substitution. Muscle alignment was plotted in Jalview version 2.11.2.6 (Waterhouse et al. 2009).The MEME Suite version 5.5.1 (Bailey et al. 2015) was used to identify conserved motifs inside OFP sequences with 5 maximum number of motifs and motif width between 3 and 80.

### 4.2. Plant material

Flower buds in stage C and fruits in stage H of the Baggiolini scale (Baggiolini 1952) were collected from two peach genotypes: the ‘UFO-4’ (UFO) flat variety and its round shape sport mutant, UFO-4Mut (MUT), which differed in a 6 Mb region at the end of the chromosome 6 due to a chromosome break and a non-homologous recombination, which lead to a homozygous region (López-Girona et al. 2017). Seven flower buds were collected and pooled from three UFO trees and five MUT trees, yielding in total. Three fruits in stage S1 were collected and pooled from three UFO trees and three MUT trees, yielding six samples in total.

Three apple varieties with contrasting fruit shapes were selected from the Spanish copy of the Apple REFPOP collection (Jung et al. 2022), as in (Dujak Riquelme 2023): the flat ‘Grand’mere’ (Gra, MUNQ 653, syn. ‘Grosse de Saint-Clément’), the round ‘Kansas Queen’ (Kan, MUNQ 565) and the oblong ‘Skovfoged’ (Sko, MUNQ 345). Three biological replicates of fruit samples were collected for each of the three varieties at 13, 61 and 98 days after anthesis (DAA). Height, width, weight, and fruit shape index (FSI) measurements were obtained at each collection point (Table S2). Height and width measures were obtained with the Tomato Analyzer software v.3 (Brewer et al. 2006). Fruit shape index (FSI) was measured as the ratio between fruit height and width (Dujak et al. 2024; Dujak Riquelme 2023).

All buds and fruit samples were frozen in liquid nitrogen and stored at -80°C until further processing for total RNA extraction.

### 4.3. RNA extraction and cDNA preparation

Total RNA from all samples was extracted with the Maxwell® RSC instrument. For the apple samples, the kit used was the Maxwell® RSC simplyRNA tissue kit (Promega) as in (Dujak Riquelme 2023; Dujak et al. 2024), while for the peach ones the kit used was the Maxwell RSC Plant RNA Kit (Promega). In both cases, the RNA was further DNase-treated with the TURBO DNA-free Kit (Invitrogen). All samples, including biological and technical replicates, were sequenced with Illumina NovaSeq 6000 PE150 Sequencing System.

### 4.4. RNAseq data analysis

The mRNA sequencing libraries were filtered by sequence quality (reads with a Phred score < 30 were removed), and remaining Illumina sequencing adaptors were trimmed using Trim-Galore version 0.6.1 (Krueger 2012). Quality control reports were obtained before and after filtering and mapping with FastQC version 0.11.5 (Andrews 2010) to ensure high-quality standards for downstream analyses. High-quality RNA sequenced libraries (RNAseq) from *M. domestica* were mapped to the apple anther-derived homozygous genotype ‘Hanfu’ (‘Dongguang’ × ‘Fuji’) genome HFTH1 (Zhang et al. 2019) with HISAT version 2.1.0 (Kim et al. 2015) using the default settings parameters. The reference genome was previously indexed with Bowtie version 2.3.4.1 (Langmead and Salzberg 2012). High-quality RNA sequenced libraries from *P. persica* were mapped to the doubled haploid cultivar ‘Lovell’ (PLOV2-2N) genotype v2.0.a1 (Verde et al. 2013; 2017) using HISAT2 version 2.2.1 (Kim et al. 2019) with default settings parameters. The reference genome was previously indexed with ‘hisat2-build’ function from HISAT2 software. Samtools version 1.9 (Li and Durbin 2009) was used to transform, index, and sort the files generated by the mappers according to the protocol’s needs.

Mapped reads were quantified and a count matrix was constructed with ‘featureCounts’ (Liao et al. 2014) function of “Rsubread” package in R. Parameters were set to paired-end sequencing, chimeric count fragments (those fragments that have their two ends aligned to different chromosomes) were excluded for apple RNAseq but allowed in peach RNAseqs. Feature type was specified as exon for apple RNAseq and transcript for peach RNAseq, annotating them at the transcript level and allowing overlapping features for the differential use during alternative splicing.

For differential expression analysis, samples were normalized using the TMM method, and statistical values were calculated with the “EdgeR” package in R (McCarthy et al. 2012; Robinson et al. 2010). In the peach RNAseq dataset, differentially expressed genes (DEGs) were identified between varieties at each developmental stage (UFO *vs* MUT in fruits and flower buds), guided by the clear stage separation observed in the PCA (Figure S3A). In the apple RNAseq dataset, comparisons were performed both across developmental stages within each variety (98 *vs* 13, 61 *vs* 13 and 98 *vs* 61 DAA in Gra, Kan and Sko), and across varieties at each developmental stages point (Kan *vs* Gra, Sko *vs* Gra and Sko *vs* Kan, at 13, 61 and 98 DAA). DEGs were filtered using an adjusted *p*-value (FDR) < 0.05 and |logFC| > 1 for peach pairwise comparisons, and FDR < 0.05 with |logFC| > 2 for apple pairwise comparisons.

Gene Ontology Enrichment Analysis (GOEA) were performed with the “gprofiler2” package in R (Kolberg et al. 2020) for *P.persica* and “TopGO” and “GO.db” packages for *M. domestica*, along with “Mdomestica_HFTH1_v1_genes2Go.xlsx” annotation file from GDR for GO term relation with gene annotation of ‘Hanfu’ HFTH1 genome. KEGG pathway enrichment were performed with the “gprofiler2” package of R for *P.persica* and “clusterProfiler” package for *M.domestica*, along with “Mdomestica_HFTH1_v1_KEGG-pathways.xlsx” annotation file from GDR for KEGG pathway relation with gene annotation of ‘Hanfu’ HFTH1 genome.

Co-expressed gene groups in apple and peach datasets were inferred using Weighted Gene Co-Expression Network Analysis (WGCNA) using the “WGCNA” R package (Langfelder and Horvath 2008; Zhang and Horvath 2005). Raw transcriptomic counts from both species were normalized to counts per million (CPM), log2-transformed, and filtered to remove genes with fewer than 10 reads in at least the 90% of samples. After filtering, 25,565 unique genes in apple fruit, 21,856 in peach flower buds, and 20,507 in peach fruits were retained for WGCNA. For the apple dataset, trait information included numerical measurements (fruit weight, height, width, and fruit shape index (FSI)) as well as categorical variables (shapes: oblong, round, flat; and developmental stage: 13, 61 and 98 DAA). For the peach dataset, trait information consisted solely of categorical data regarding shape (UFO and MUT). Unsigned networks were built manually by transforming adjacency matrices into Topological Overlap Matrix (TOM). Soft-thresholding powers of 8 (apple) and 9 (peach) were defined using a minimum module size of 30 genes, a deep split parameter of 2, and a branch merge cut height of 0.20 for all datasets, except for peach fruit, where a cut height of 0.10 was used. In the apple dataset, given the large number of module-trait correlations, *p*-values were adjusted using the false discovery rate (FDR) method (Benjamini and Hochberg 1995).

Hierarchical clustering using the complete method of hclust function in R was carried out to identify 9 clusters of expression profiles within each module identified by the WGCNA.

## CRediT authorship contribution statement

M.J.A. supervised the study. C.D. and A.F. conducted the laboratory experiments. V.C.A. and B.E.G.C. processed the sequenced libraries. V.C.A. carried out phylogenetic, transcriptomic and network analysis. V.C.A. and M.J.A. wrote the manuscript. All authors reviewed the manuscript.

## Acknowledgments

VCA was recipient of grant PRE2019-088780 funded by MICIU/AEI/ 10.13039/501100011033 and by “ESF Investing in your future”. A.F. was recipient of grant BES-2016-079060 funded by MICIU/AEI/10.13039/501100011033 and by “ESF Investing in your future”. CD was supported by “DON CARLOS ANTONIO LOPEZ” Abroad Postgraduate Scholarship Program, BECAL-Paraguay.

This research was supported by grants RTI2018-100795-B-I00 funded by MICIU/AEI/10.13039/501100011033 and by “ERDF A way of making Europe”; PID2021-128885OB-I00, and PID2024-163074OB-I00 funded by MICIU/AEI/10.13039/501100011033 and by “ERDF/EU”; This work was also supported by grants SEV-2015-0533 and CEX2019-000902-S funded by MICIU/AEI/10.13039/501100011033; by the CERCA Programme/Generalitat de Catalunya; and by the SGR Program (2021-SGR-00756) funded by Direcció General de Recerca (DGR) del Departament de Recerca i Universitats (REU).

## Supplementary Figures and tables

**Table S1: OFP protein sequences used for phylogenetic analysis**

**Table S2: Morphometric measurements (weight, height, width, and FSI) of apple samples**

**Table S3: Differentially expressed genes (DEGs) identified in peach samples**

**Table S4: Differentially expressed genes (DEGs) identified in apple samples**

**Table S5: DEGs from peach fruit RNAseq data within each WGCNA module and cluster**

**Table S6: DEGs from peach flower bud RNAseq data within each WGCNA module and cluster**

**Table S7: DEGs from apple fruit RNAseq data within each WGCNA module and cluster**

**Figure S1. Multiple alignment of 127 OFP amino acid sequences.** The OVATE domain region is highlighted within a grey dashed rectangle. OFPs lacking the OVATE domain are marked in black and framed in a red rectangle. Identical residues within the OVATE domain are shaded in dark blue; similar residues are shaded in light blue.

**Figure S2. Conserved motif analysis of 127 OFPs.** Motifs identified using MEME are color-coded in the Motif Locations column. Sequences are sorted and included in numbered and colored rectangles as in the clades of the phylogenetic tree in Figure 1.

**Figure S3. Transcriptomic analysis of fruit shape and developmental stages in *Prunus persica*.** (A) Principal component analysis (PCA) of peach RNAseq samples. (B) Venn diagrams with the number of DEGs found in the shape pairwise comparison across developmental stages.

**Figure S4. Transcriptomic analysis of fruit shape and developmental stages of *Malus domestica*.** Principal component analysis (PCA) of apple RNAseq samples in dimensions 1-2 (A), and 2-3 (B). (C-E) Venn diagrams showing DEGs identified in pairwise comparisons.

**Figure S5. GO enrichment analysis of developmental stage comparisons across fruit shapes in apple.** BP: Biological Process, CC: Cellular Component, MF: Molecular Function, KEGG: Kyoto Encyclopaedia of Genes and Genomes (KEGG) pathway.

**Figure S6. Plant Hormone Signal Transduction (KEGG: ko04075) map highlighting deregulated genes of developmental stages comparison in each apple variety.** (A) Gra_61vs13, (B) Gra_98vs13, (C) Gra_98vs61, (D) Kan_61vs13, (E) Kan_98vs13, (F) Kan_98vs61, (G) Sko_61vs13, (H) Sko_98vs13, (I) Sko_98vs61. Highlighted genes are colored by the logFC in red for downregulation and in blue for upregulation.

**Figure S7. GO enrichment analysis of shape comparisons across developmental stages in apple.** BP: Biological Process, CC: Cellular Component, MF: Molecular Function, KEGG: Kyoto Encyclopaedia of Genes and Genomes (KEGG) pathway.

**Figure S8. WGCNA of peach fruit transcriptomic data.** (A) Dendrogram and phenotypic information of RNA-seq samples. (B) Hierarchical clustering of co-expressed gene modules. (C) Heatmap of gene expression in the “palevioletred” module. (D) Line plots of clustered gene expression patterns within the module.

**Figure S9. WGCNA of peach flower bud transcriptomic data.** (A) Dendrogram and phenotypic information of RNA-seq samples. (B) Hierarchical clustering of co-expressed gene modules. (C) Heatmap of gene expression in the “aquamarine4” module. (D) Line plots of clustered gene expression patterns within the module.

**Figure S10. WGCNA of apple fruit transcriptomic data.** (A) Dendrogram and phenotypic information of RNA-seq samples. (B) Hierarchical clustering of co-expressed gene modules. (C, E, G) Heatmaps of gene expression in “pink,” “red,” and “green” modules. (D, F, H) Line plots of clustered gene expression patterns within each module.

