## Supplementary material for "Crosstalk between Ovate Family Proteins, plant hormones, and microtubule dynamics regulating fruit shape": Figure S1-S10

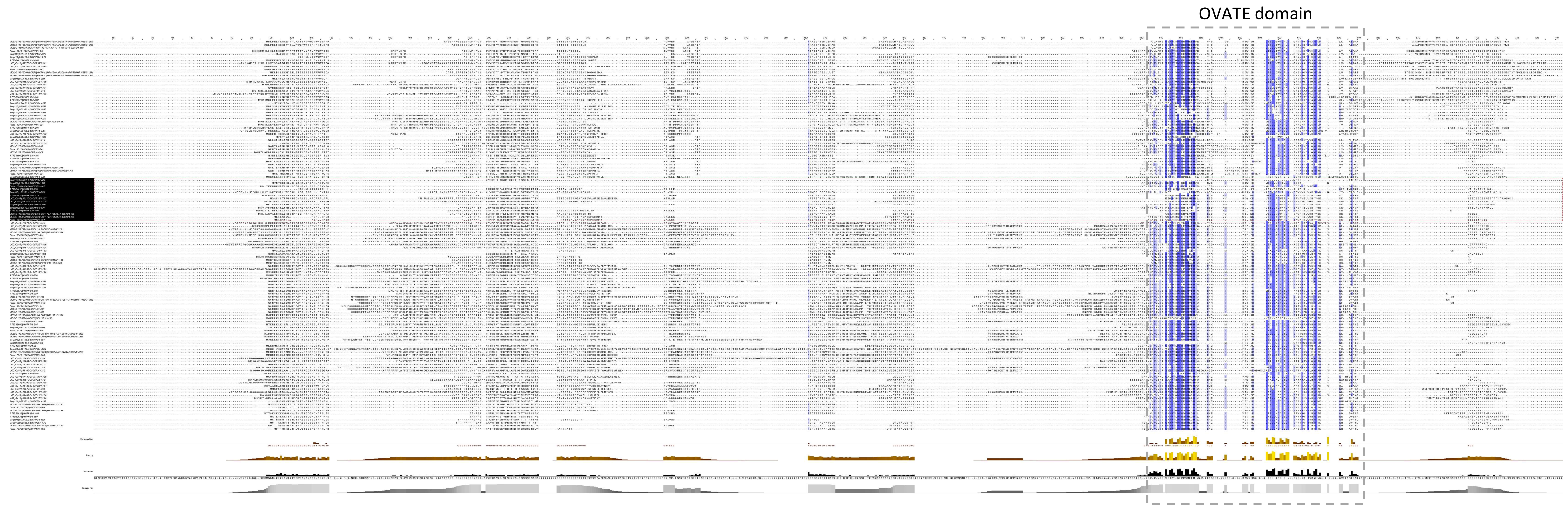


Figure S1. Multiple sequence alignment of 127 OFP amino acid sequence. The OVATE domain region is highlighted with a grey-dashed rectangle. OFPs lacking the OVATE domain are marked in black and framed in a red rectangle. Identical residues are shaded in dark blue; similar residues are shaded in light blue.

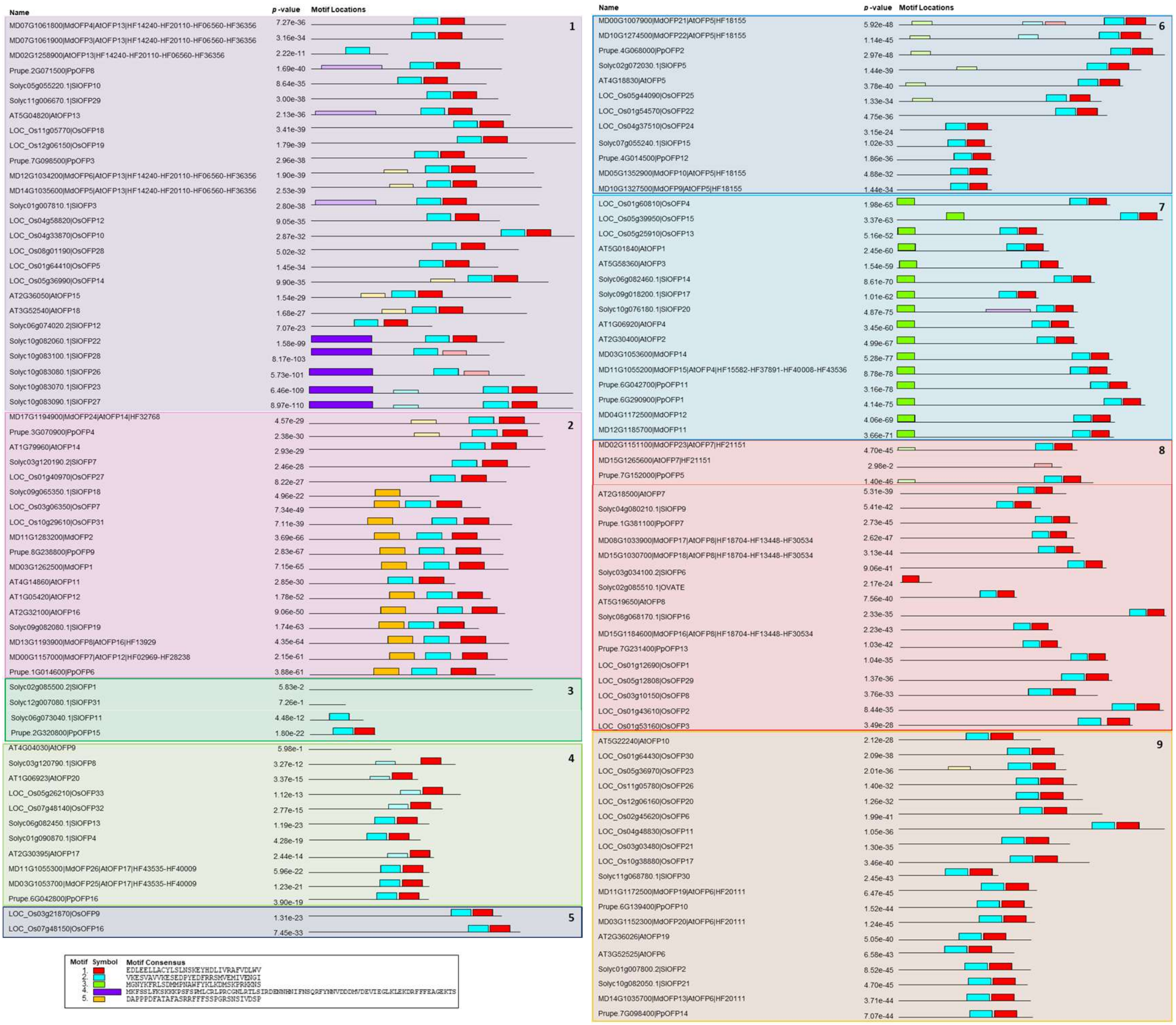

Figure S2. Conserved motif analysis of 127 OFPs. Motifs identified using MEME are color-coded in the Motif Locations column. Sequences are sorted and included in numbered and colored rectangles as in the clades of the phylogenetic tree in Figure 1.

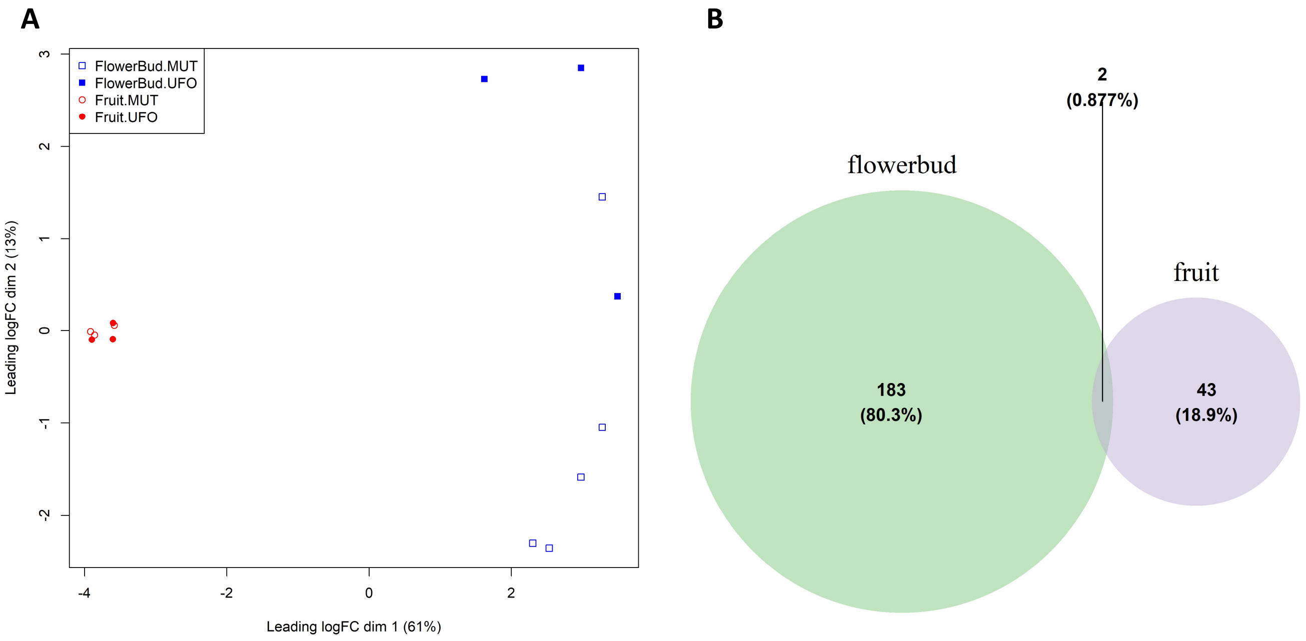


Figure S3. Transcriptomic analysis of fruit shape and developmental stages in Prunus persica. (A) Principal component analysis (PCA) of peach RNAseq samples. (B) Venn diagrams with the number of DEGs found in the shape pairwise comparison across developmental stages.

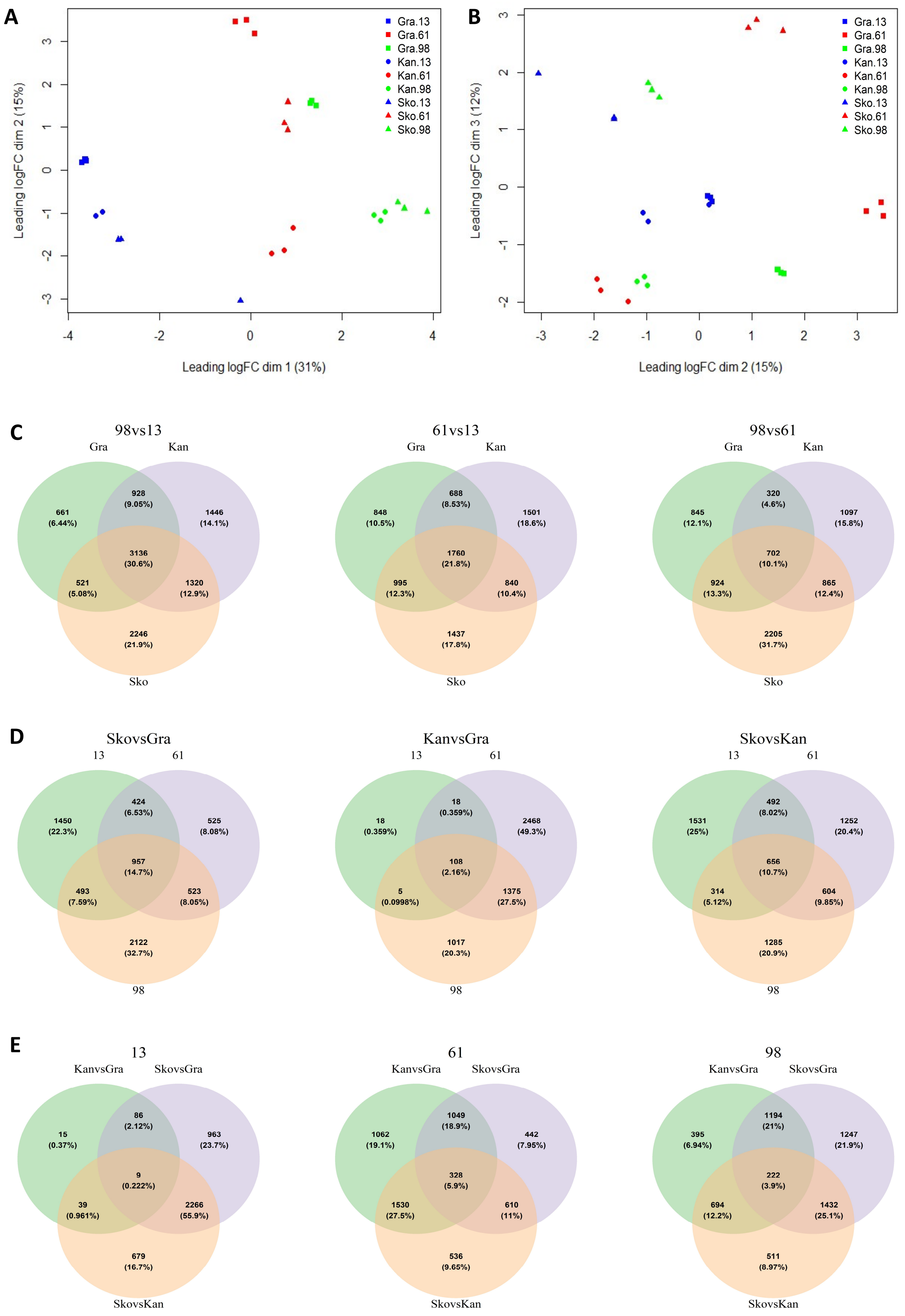


Figure S4. Transcriptomic analysis of fruit shape and developmental stages of Malus domestica. Principal component analysis (PCA) of apple RNAseq samples in dimensions 1-2 (A), and 2-3 (B). (C-E) Venn diagrams showing DEGs identified in pairwise comparisons.

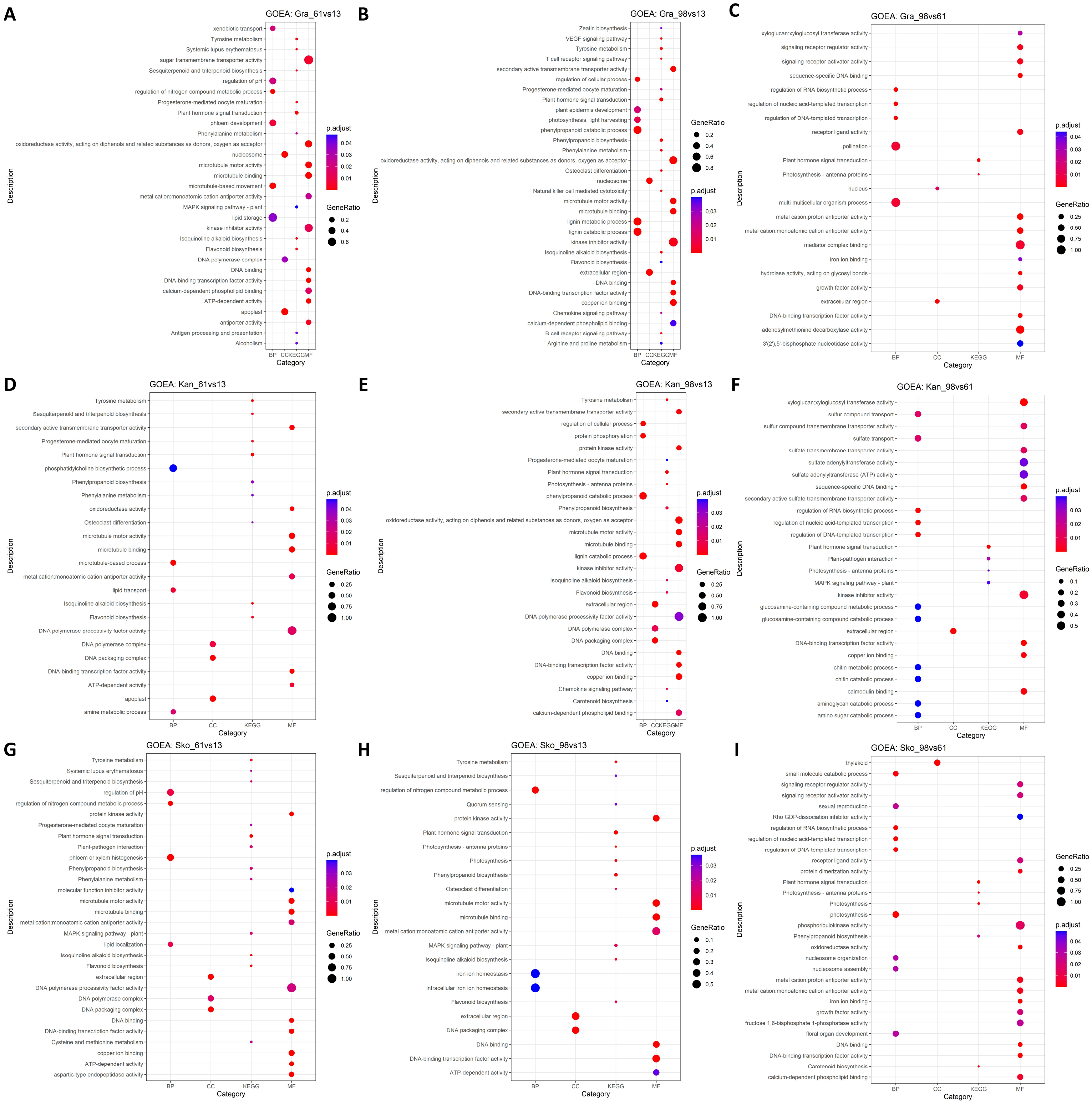


Figure S5. GO enrichment analysis of developmental stage comparisons across fruit shapes in apple. BP: Biological Process, CC: Cellular Component, MF: Molecular Function, KEGG: Kyoto Encyclopaedia of Genes and Genomes (KEGG) pathway.

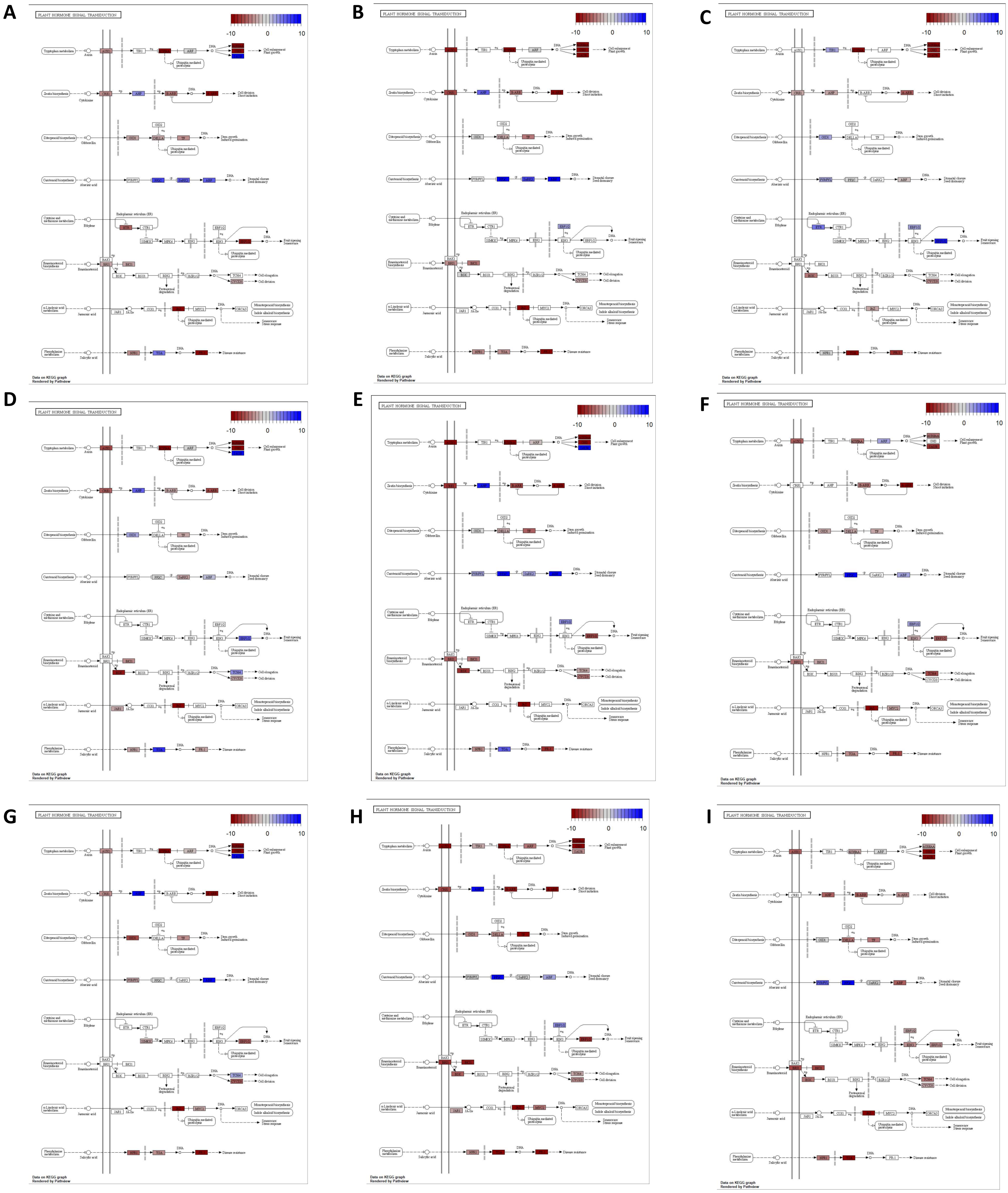


Figure S6. Plant Hormone Signal Transduction (KEGG: ko04075) map highlighting deregulated genes of developmental stages comparison in each apple variety (A) Gra_61vs13, (B) Gra_98vs13, (C) Gra_98vs61, (D) Kan_61vs13, (E) Kan_98vs13, (F) Kan_98vs61, (G) Sko_61vs13, (H) Skoa_98vs13, (I) Sko_98vs61. Highlighted genes are colored by the logFC in red for downregulation and in blue for upregulation.

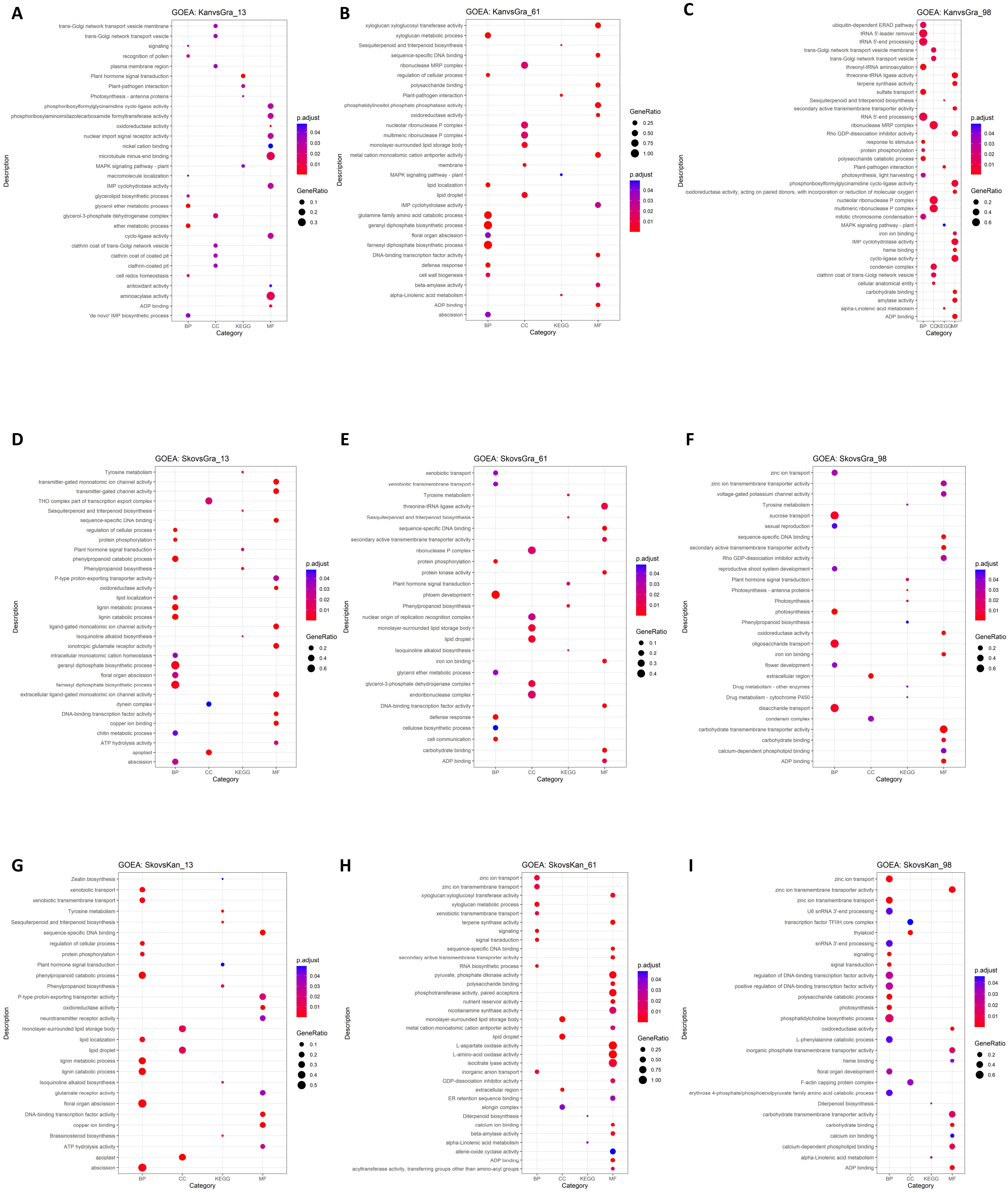


Figure S7. GO enrichment analysis of shape comparisons across developmental stages in apple. BP: Biological Process, CC: Cellular Component, MF: Molecular Function, KEGG: Kyoto Encyclopaedia of Genes and Genomes (KEGG) pathway.

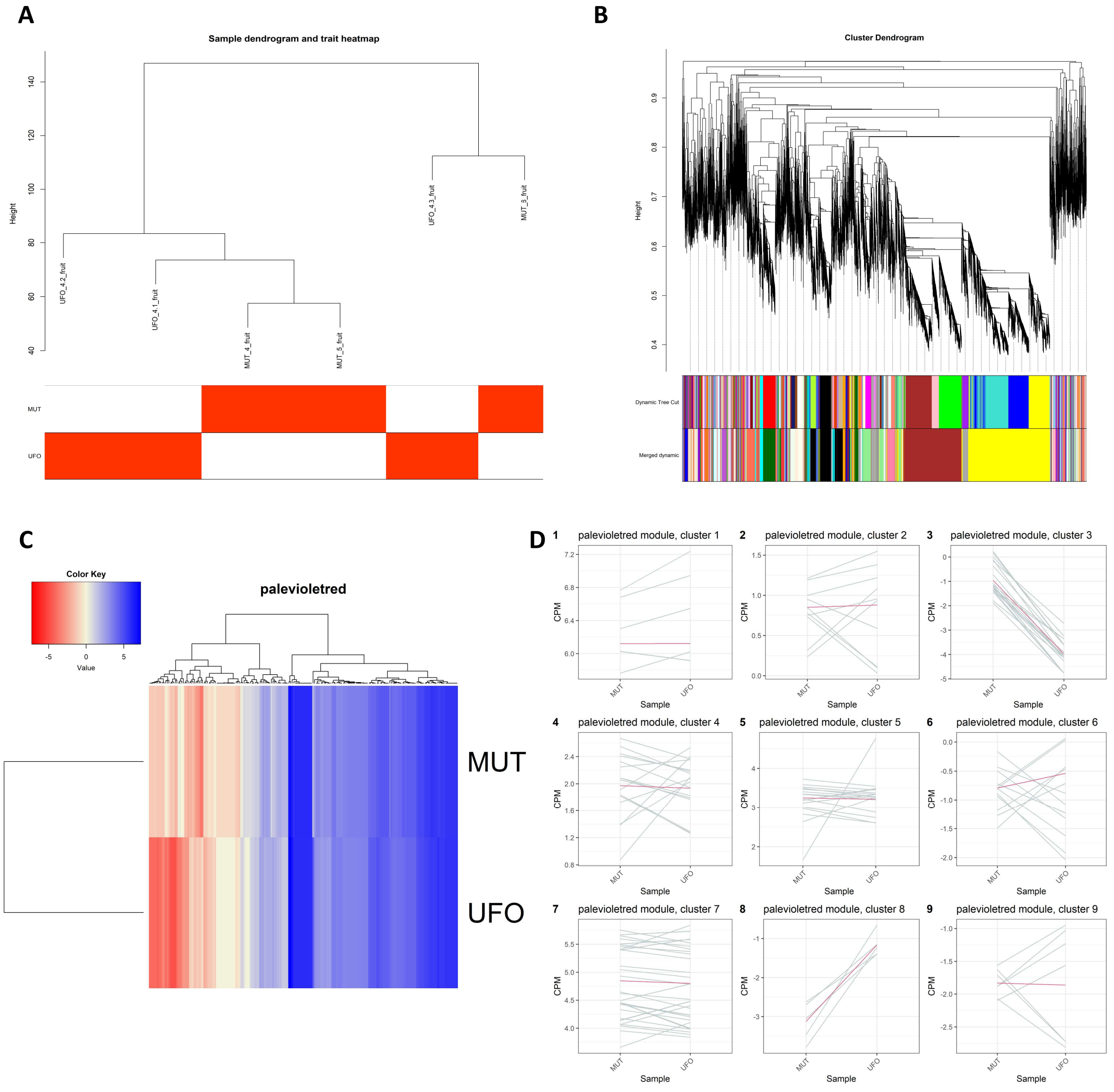


Figure S8. WGCNA of peach fruit transcriptomic data. (A) Dendrogram and phenotypic information of RNA-seq samples. (B) Hierarchical clustering of co-expressed gene modules. (C) Heatmap of gene expression in the “palevioletred” module. (D) Line plots of clustered gene expression patterns within the module.

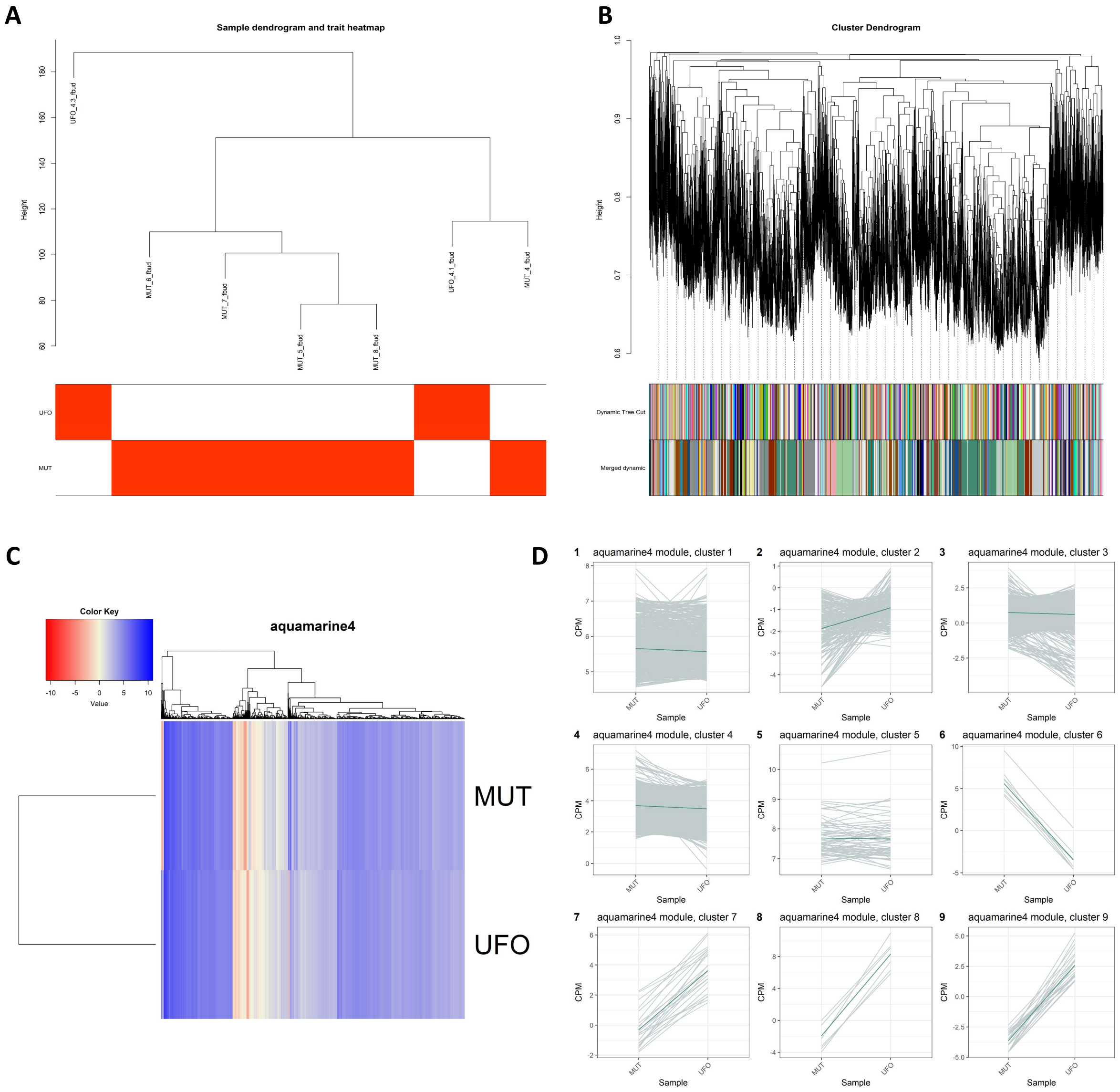


Figure S9. WGCNA of peach flower bud transcriptomic data. (A) Dendrogram and phenotypic information of RNA-seq samples. (B) Hierarchical clustering of co-expressed gene modules. (C) Heatmap of gene expression in the “aquamarine4” module. (D) Line plots of clustered gene expression patterns within the module.

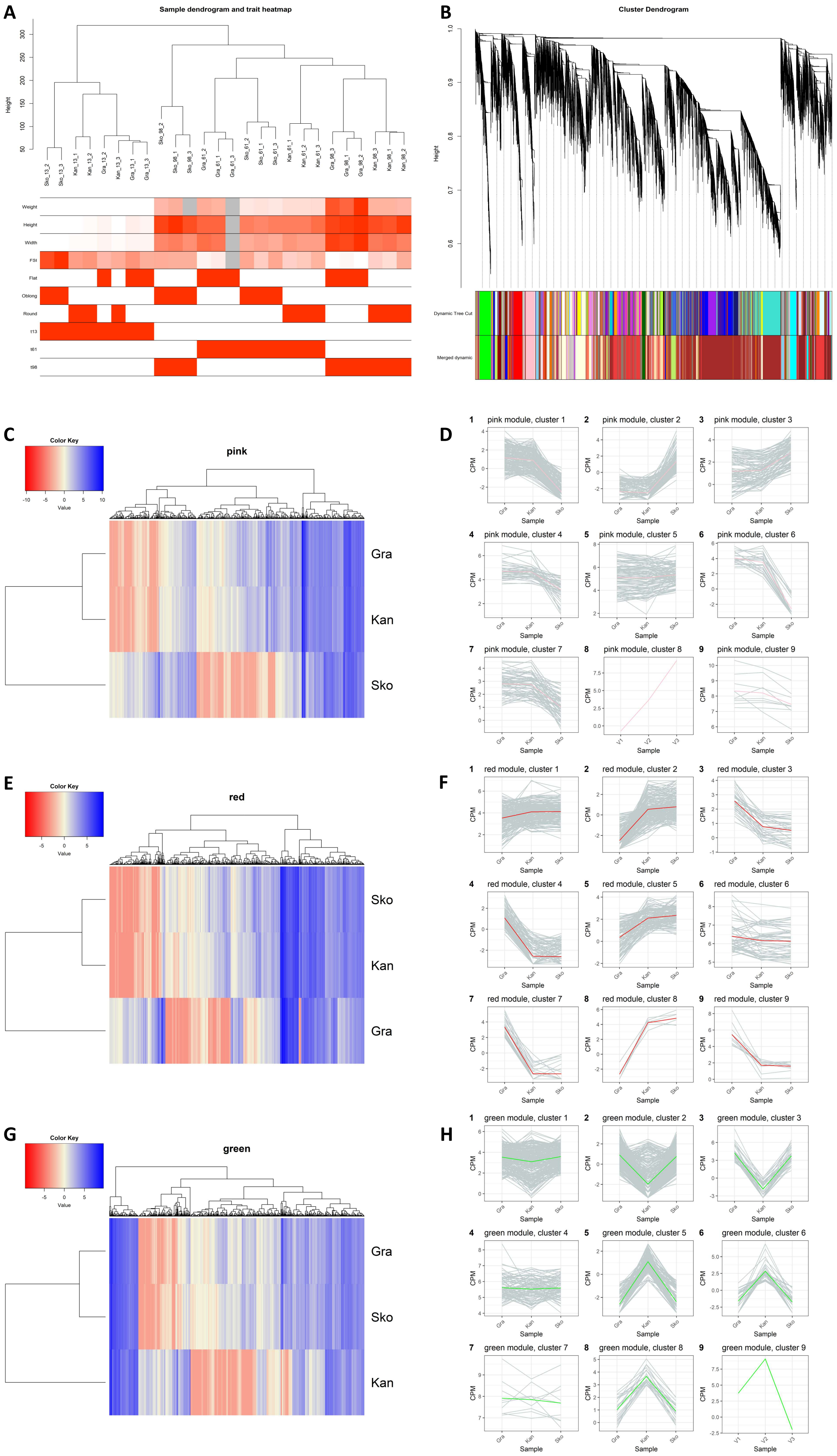
A
Sample dendrogram and trait heatmap
Height
Sko_13_2
Sko_13_3
Kan_13_1
Kan_13_2
Gra_13_2
Kan_13_3
Gra_13_1
Gra_13_3
Sko_98_2
Sko_98_1
Sko_98_3
Gra_61_2
Gra_61_1
Gra_61_3
Sko_61_2
Sko_61_1
Sko_61_3
Kan_61_1
Kan_61_2
Kan_61_3
Gra_98_3
Gra_98_1
Gra_98_2
Kan_98_3
Kan_98_1
Kan_98_2
Weight
Height
Width
FSI
Flat
Oblong
Round
t13
t61
t98
B
Cluster Dendrogram
Height
Dynamic Tree Cut
Merged dynamic
C
Color Key
Value
pink
Gra
Kan
Sko
D
1 pink module, cluster 1
2 pink module, cluster 2
3 pink module, cluster 3
4 pink module, cluster 4
5 pink module, cluster 5
6 pink module, cluster 6
7 pink module, cluster 7
8 pink module, cluster 8
9 pink module, cluster 9
CPM
Sample
E
Color Key
Value
red
Sko
Kan
Gra
F
1 red module, cluster 1
2 red module, cluster 2
3 red module, cluster 3
4 red module, cluster 4
5 red module, cluster 5
6 red module, cluster 6
7 red module, cluster 7
8 red module, cluster 8
9 red module, cluster 9
CPM
Sample
G
Color Key
Value
green
Gra
Sko
Kan
H
1 green module, cluster 1
2 green module, cluster 2
3 green module, cluster 3
4 green module, cluster 4
5 green module, cluster 5
6 green module, cluster 6
7 green module, cluster 7
8 green module, cluster 8
9 green module, cluster 9
CPM
Sample

Figure S10. WGCNA of apple fruit transcriptomic data. (A) Dendrogram and phenotypic information of RNA-seq samples. (B) Hierarchical clustering of co-expressed gene modules. (C, E, G) Heatmaps of gene expression in “pink,” “red,” and “green” modules. (D, F, H) Line plots of clustered gene expression patterns within each module.
